# Lignin impairs Cel7A degradation of cellulose by impeding enzyme movement and not by acting as a sink

**DOI:** 10.1101/2023.08.23.554521

**Authors:** Zachary K. Haviland, Daguan Nong, Nerya Zexer, Ming Tien, Charles T. Anderson, William O. Hancock

**Author notes:** **Author Contributions:** Z.K.H, D.N, N.Z., M.T., C.T.A. and W.O.H. designed research; Z.K.H. and D.G. performed research; Z.K.H. and D.G. analyzed data; Z.K.H., C.T.A. and W.O.H. wrote the paper. **Competing Interest Statement:** The authors declare that they have no known competing financial interests or personal relationships that could have appeared to influence the work reported in this paper.

## Abstract

**Background:** Cellulose degradation by cellulases has been studied for decades due to the potential of using lignocellulosic biomass as a sustainable source of bioethanol. In plant cell walls, cellulose is bonded together and strengthened by the polyphenolic polymer, lignin. Because lignin is tightly linked to cellulose and is not digestible by cellulases, is thought to play a dominant role in limiting the efficient enzymatic degradation of plant biomass. Removal of lignin via pretreatments currently limits the cost-efficient production of ethanol from cellulose, motivating the need for a better understanding of how lignin inhibits cellulase-catalyzed degradation of lignocellulose. Work to date using bulk assays has suggested three possible inhibition mechanisms: lignin blocks access of the enzyme to cellulose, lignin impedes progress of the enzyme along cellulose, or lignin binds cellulases directly and acts as a sink.

**Results:** We used single-molecule fluorescence microscopy to investigate the nanoscale dynamics of Cel7A from *Trichoderma reesei*, as it binds to and moves along purified bacterial cellulose in vitro. Lignified cellulose was generated by polymerizing coniferyl alcohol onto purified bacterial cellulose, and the degree of lignin incorporation into the cellulose meshwork was analyzed by optical and electron microscopy. We found that Cel7A preferentially bound to regions of cellulose where lignin was absent, and that in regions of high lignin density, Cel7A binding was inhibited. With increasing degrees of lignification, there was a decrease in the fraction of Cel7A that moved along cellulose rather than statically binding. Furthermore, with increasing lignification, the velocity of processive Cel7A movement decreased, as did the distance that individual Cel7A molecules moved during processive runs.

**Conclusions:** In an in vitro system that mimics lignified cellulose in plant cell walls, lignin did not act as a sink to sequester Cel7A and prevent it from interacting with cellulose. Instead, lignin both blocked access of Cel7A to cellulose and impeded the processive movement of Cel7A along cellulose. This work implies that strategies for improving biofuel production efficiency should target weakening interactions between lignin and cellulose surface, and further suggest that nonspecific adsorption of Cel7A to lignin is likely not a dominant mechanism of inhibition.

## Introduction

Biofuels are a renewable energy source that can assist in the transition to a low-carbon economy as the world faces the dual challenges of climate change and increasing energy demand (1, 2). Cellulose and lignin, two of the most abundant biopolymers found in nature, are structural components of the plant cell wall that have become targets for biofuel production, with a variety of strategies being implemented to convert these two substrates into usable forms of energy (2–4). Cellulose is a homopolymer of β-1,4-linked glucose units, with parallel glucan chains bonding to form partially crystalline microfibrils (5, 6). Lignin is a polyphenolic polymer generated by radical coupling of monolignols that interacts tightly with cellulose in the cell wall (7, 8). Cellulosic biomass can be converted into biofuels after deconstruction by cellulases and fermentation of the resulting glucose (4, 9). However, enzymatic cellulose degradation is currently inefficient due to both the limited accessibility of cellulose chains when packed into a crystalline lattice, and obstruction by other cell wall components such as lignin and hemicellulose (10–13). Lignin cannot be hydrolyzed by cellulases, making plant biomass recalcitrant to enzymatic digestion, but the mechanisms by which lignin impedes cellulase activity are poorly understood (14–16). In current schemes for biofuel production, lignin is removed from cellulosic biomass by thermochemical and acidic pretreatments that are costly and reduce the feasibility, scalability, and sustainability of the process (17–19). Thus, understanding how lignin inhibits the degradation of cellulose by cellulases is crucial for optimizing biofuel production from lignocellulose.

A model cellulase used to study cellulose degradation is the cellobiohydrolase I, Cel7A. *Tr*Cel7A is a processive exoglucanase from *Trichoderma reesei* (teleomorph *Hypocrea jecorina*) that binds to the reducing end of cellulose chains. Its catalytic domain hydrolyzes the glycosidic bonds in cellulose, releasing the disaccharide cellobiose as a product, while the cellulose binding module is thought to bind to crystalline cellulose and thus enhance enzyme affinity (6, 20, 21). Removal of lignin from plant-derived lignocellulose improves the rate of cellulose degradation in bulk assays (13, 16, 22). Similarly, polymerizing lignin onto isolated cellulose *in vitro* decreases the bulk cellulase activity (22–25). These and other prior studies have posited three potential mechanisms by which lignin inhibits cellulose degradability: 1) lignin might physically block the initial binding of Cel7A to the cellulose surface (13, 26); 2) lignin might impede the processive catalysis of Cel7A bound to cellulose, effectively acting as a roadblock (27); 3) lignin might act as a “sink” for Cel7A by irreversibly adsorbing the enzyme (Fig. 1) (25, 28–30). Thus, although there is general agreement that lignin inhibits cellulase activity, the precise mechanism of inhibition and the best strategy for relieving this inhibition are not clear.

**Figure 1:**
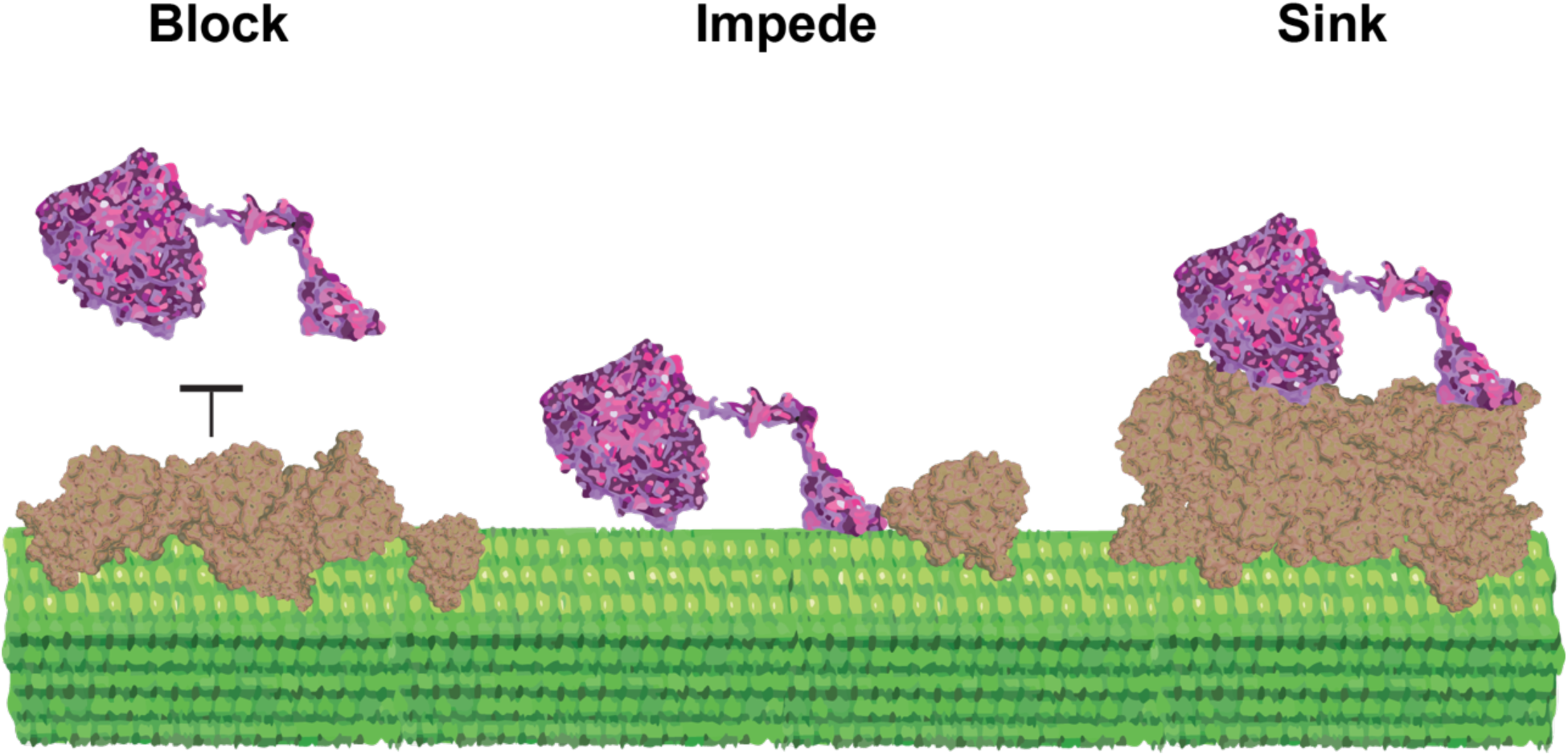
Potential inhibition mechanisms. Lignin may inhibit Cel7A on cellulose by blocking binding of enzymes to cellulose, by impeding the progress of enzymes bound to cellulose, or by acting as a sink by nonspecifically adsorbing enzymes.

One shortcoming in the field is that most prior analyses of cellulase inhibition by lignin lack direct observations of the interaction between lignin and cellulases (13, 16, 22, 24, 25). However, direct visualization of cellulase enzymes is possible using single-molecule microscopy (27, 31). We have constructed a high-resolution microscope that can track, at nanometer resolution, thousands of individual Cel7A molecules performing cellulose degradation on immobilized cellulose (32). We found that Cel7A reversibly transitions between free diffusion in solution, static binding to cellulose, and processive movement along cellulose (33). Here, we extended this single-molecule fluorescence approach to test potential molecular mechanisms by which lignin inhibits cellulose degradation by Cel7A. Synthetic analogs of plant cell walls were constructed by polymerizing G-lignin *in vitro* onto bacterial cellulose at varying lignin-to-cellulose ratios (34). This approach eliminates potential effects of other wall components like hemicellulose and pectin, allowing us to focus only on lignin bound to cellulose. The resulting lignified cellulose also serves as a model of pretreated native lignocellose that is used industrially. We asked the following three questions: 1) Does lignin deposition on cellulose inhibit Cel7A binding to the cellulose surface? 2) Does lignin alter Cel7A dynamics during processive movement along cellulose? 3) Does Cel7A accumulate on lignin due to irreversible binding?

Using a combination of scanning electron microscopy (SEM) and interference reflection microscopy (IRM), we found that *in vitro*-polymerized lignin formed structures that enclose cellulose strands and that Cel7A binding to lignified cellulose was negatively correlated with lignin. Using total internal reflection fluorescence microscopy (TIRFM), we quantified the dynamics of quantum dot-labeled Cel7A molecules and found that lignin decreased the fraction of moving enzymes, as well as their run lengths and velocities. These data are inconsistent with lignin acting as a sink that irreversibly binds Cel7A. Instead, the data indicate that lignin inhibits Cel7A by blocking its initial interaction with the cellulose surface and by impeding the progress of moving Cel7A molecules.

## Results

### *In vitro* polymerized lignin occludes the surface of cellulose

To examine how lignin alters cellulose degradation by Cel7A, we created lignified cellulose by depositing lignin onto *Acetobacter* cellulose *in vitro* using a mixture of coniferyl alcohol, hydrogen peroxide and horseradish peroxidase (34). To vary the degree of lignification, the reaction contained 4.25 mM cellulose and between 0.11 mM and 9 mM coniferyl alcohol and hydrogen peroxide. Hereafter, the CA concentrations added to the reactions are used to differentiate the lignin samples. To assess the relative coverage of the cellulose by lignin, we first examined samples by interference reflection microscopy (IRM) (35, 36). Flow cells were created using a plasma-cleaned coverslip and a microscope slide, and the lignocellulose samples were dried in the flow cell to adhere to the coverslip before being rehydrated with buffer. Under IRM, the cellulose-only sample showed large tangles of cellulose strands with the strands becoming sparser around the edges of the cellulose masses, allowing smaller cellulose bundles to be visualized (Fig. 2A). In contrast, in the lignin-only sample, large aggregates of lignin were visible that had much more contrast than the cellulose samples and had a smooth morphology under IRM compared to the rough surface of the cellulose (Fig. 2B). By IRM, the 0.11 mM and 0.33 mM CA samples were indistinguishable from the cellulose-only controls (Fig. 2C and Fig. S1D), but at concentrations of 1 mM CA and higher, the lignin was visible under IRM as small round aggregates dispersed across the cellulose surface (Fig. 2D). At the highest CA concentration of 9 mM, large patches of lignin covered significant portions of the cellulose surface and appeared to be integrated between the cellulose strands (Fig. 2E).

**Figure 2:**
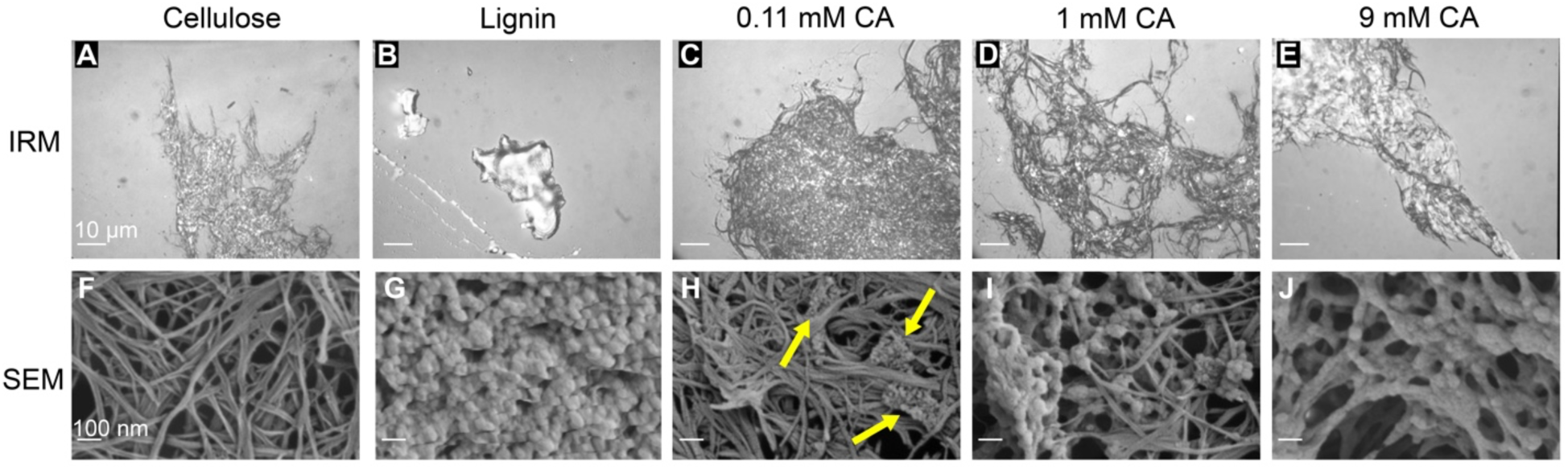
Morphology of lignin polymerized with acetobacter cellulose *in vitro*. First column: cellulose only; second column: lignin only; third through fifth columns: cellulose with lignin polymerized from increasing concentrations of coniferyl alcohol (CA). Top row are images from interference reflection microscopy (IRM), scale bars = 10 µm. Bottom row are images from scanning electron microscopy (SEM), scale bars = 100 nm. An additional image with lignin highlighted in false colored is provide in Fig. S2.

Next, we used scanning electron microscopy (SEM) to probe the nanostructure of the synthetic lignocellulose. The cellulose-only sample showed fibrils of various widths and lengths that formed an intricate meshwork with varying depths (Fig. 2F). In contrast, the lignin-only sample appeared as a mass of globular structures that created a rough surface with no clear organization (Fig. 2G). In the 0.11 mM CA sample, nanoscale cauliflower-like lignin aggregates were intertwined in the cellulose strands as indicated by yellow arrows in Figure 2H; increasing aggregate densities were observed in the 1 mM sample with some aggregates on the cellulose surface and others entangled in the cellulose meshwork (Fig. 2I). In the 9 mM CA sample, some lignin patches encased regions of cellulose, and in some instances appeared to constrict multiple cellulose strands into larger bundles (Fig. 2J). It should be noted that the deposition of lignin onto cellulose was heterogenous, with some areas of cellulose having more lignin than other areas in the same sample (Fig. S2). Together, these results establish that increasing concentrations of CA result in both greater amounts of lignin deposition on cellulose and different lignocellulose morphologies.

### Cel7A does not preferentially accumulate on *in vitro* polymerized lignin

The first question we addressed was the degree to which Cel7A directly interacts with lignin. On one hand, lignin might physically block binding of Cel7A to cellulose (Fig. 1, left) (13, 26). On the other hand, lignin may act as a “sink” by nonspecifically binding Cel7A and thus indirectly preventing the enzyme from interacting with cellulose (Fig. 1, right) (25, 28–30). To test these competing hypotheses, we visualized the locations of Cel7A binding events on the immobilized cellulose and compared them to the position of the deposited lignin. We focused our analysis on the 3 mM and 9 mM samples that contained micron-scale lignin patches that were considerably larger than the ∼300 nm point-spread-function of the microscope (32). Cellulose was imaged with IRM, Cel7A labeled with quantum dots (Qdots) were introduced, and the binding events on lignocellulose were recorded for 500 seconds using total internal reflection fluorescence microscopy (TIRFM) (33). Following a washout of the Cel7A with buffer, the lignin-specific dye Basic Fuchsin (37) was washed in and the location of the lignin was imaged by TIRFM. Finally, the image of fluorescently labeled lignin was overlayed with a maximum projection of the Cel7A video to show the binding locations for all the molecules in each video (Fig. 3).

**Figure 3:**
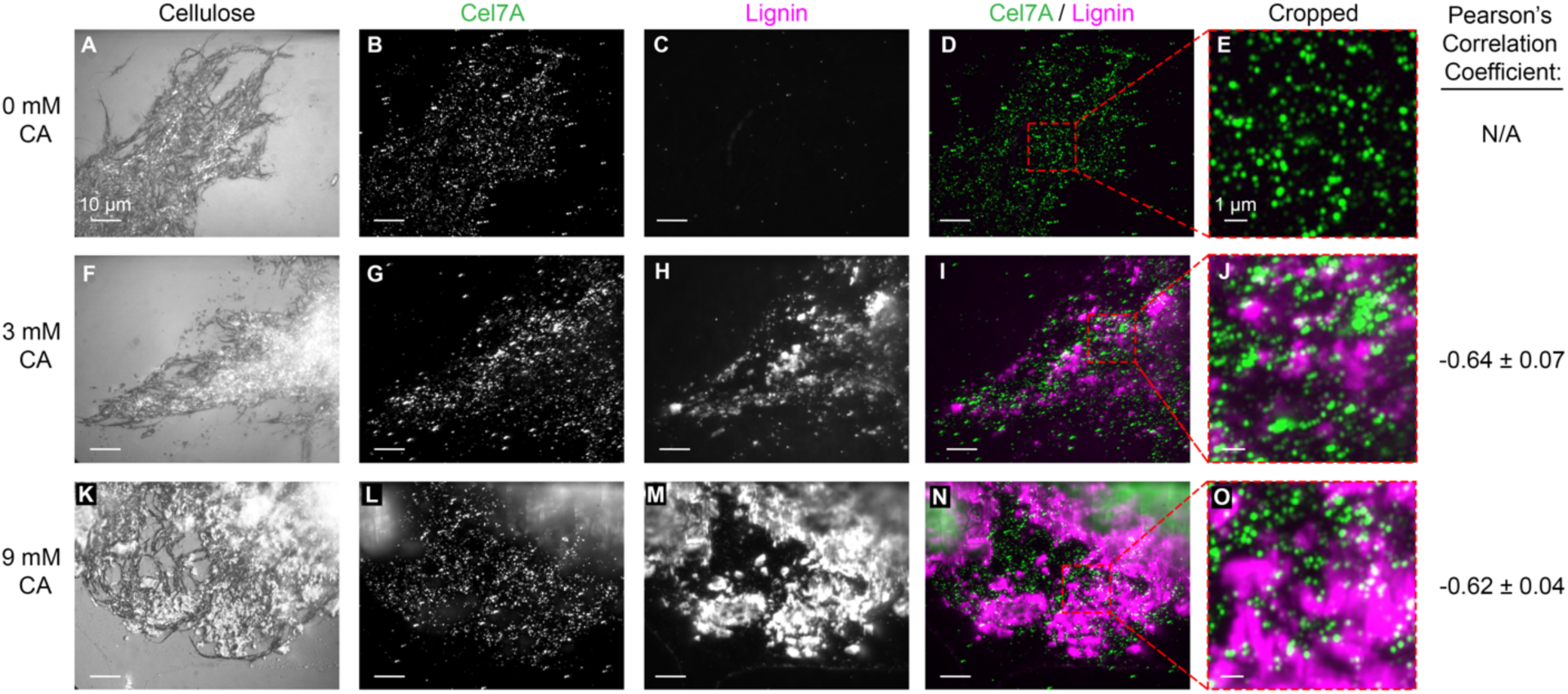
Cel7A enzymes bind to immobilized cellulose but not to lignin. First column: interference reflection microscopy images of synthetic lignocellulose immobilized on a glass coverslip. Second column: total internal reflection fluorescence microscopy images of Qdot-labeled Cel7A as they bind to cellulose over 500 seconds. Third column: lignin fluorescently stained by the lignin dye Basic Fuchsin. Fourth column: overlay of Qdot-labeled Cel7A (green) and lignin (magenta) images. Fifth column: cropped regions of the overlay images shown in the fourth column. For columns 1-4: scale bar = 10 µm. For column 5: scale bar = 1 µm.

We next compared the sites of Cel7A binding with the position of the deposited lignin. In the cellulose-only samples, no visible Basic Fuchsin fluorescence was observable (Fig. 3C), and the Cel7A binding density was relatively uniform across the immobilized cellulose (Fig. 3B). In the 3 mM CA samples, small patches and puncta of lignin were visible (Fig. 3H), whereas large patches of lignin were apparent in the 9 mM CA samples (Fig. 3M). In overlay images, few Cel7A molecules appeared to co-localize with lignin (Fig. 3I and Fig. 3N). To more quantitatively assess the spatial correlation between Cel7A and lignin, we used an ImageJ plug-in that compares the fluorescent intensities of different fluorescent channels for each pixel and calculates a Pearson’s correlation coefficient (38). A “sink” model predicts a positive correlation with +1 denoting perfect co-localization, a “blocking” model predicts a negative correlation with −1 denoting perfect anticorrelation, and zero correlation is predicted if lignin has no effect on Cel7A binding to cellulose (39). An average Pearson’s correlation coefficient of −0.638 ± 0.068 (mean ± SD, N = 3 experiments) was calculated for the 3 mM CA samples and a −0.615 ± 0.037 (mean ± SD, N = 3 experiments) coefficient was calculated for the 9 mM CA samples. From these negative correlations, we conclude that with our experimental conditions in this *in vitro* environment, lignin does not act as a “sink” by nonspecifically binding Cel7A. Instead, the data are consistent with lignin blocking the binding of Cel7A to cellulose. As a control to test whether bovine serum albumin (BSA) used to block nonspecific binding from the glass surface was blocking Cel7A binding to lignin, we repeated the experiment without adding BSA to the solutions. In the absence of BSA the correlation was 0.037, indicating the Cel7A bound to a similar extent to cellulose and lignin (Fig. S3). Thus, in the absence of blocking protein, Cel7A can non-specifically bind to lignin, but lignin does not act as a preferential sink for binding Cel7A.

### *In vitro* polymerized lignin impedes Cel7A motion on cellulose

To assess how lignin alters the behavior of Cel7A on cellulose, single-molecule fluorescence imaging was used to track quantum dot-labeled Cel7A molecules binding to and moving along immobilized lignocellulose. Videos were recorded at 1 frame/s for 1000 s and the resulting trajectories of individual particles were tracked by fitting a 2D gaussian distribution to the point-spread function of the quantum dots in every frame using the program FIESTA (40). To subtract stage drift, TetraSpeck beads were non-specifically fixed to the glass surface and used as fiduciary markers (32). The resulting drift-corrected trajectories were analyzed using a custom-made MATLAB code that plots both X-Y positions and the distance traveled from the origin versus time. Many of the tracked molecules remained static, defined as a displacement of less than 10 nm from the original location over the duration of a binding event. Other molecules displayed processive segments, defined as the enzyme moving continuously for at least 5 s for a distance of at least 10 nm. A minimum velocity of 0.1 nm/s was applied to the analysis to differentiate processive movement from stage drift that was not fully corrected. Exemplary traces of X-Y positions for processive molecules, as well as plots displaying the distance from the origin over time, are shown in Figure 4A-D.

**Figure 4:**
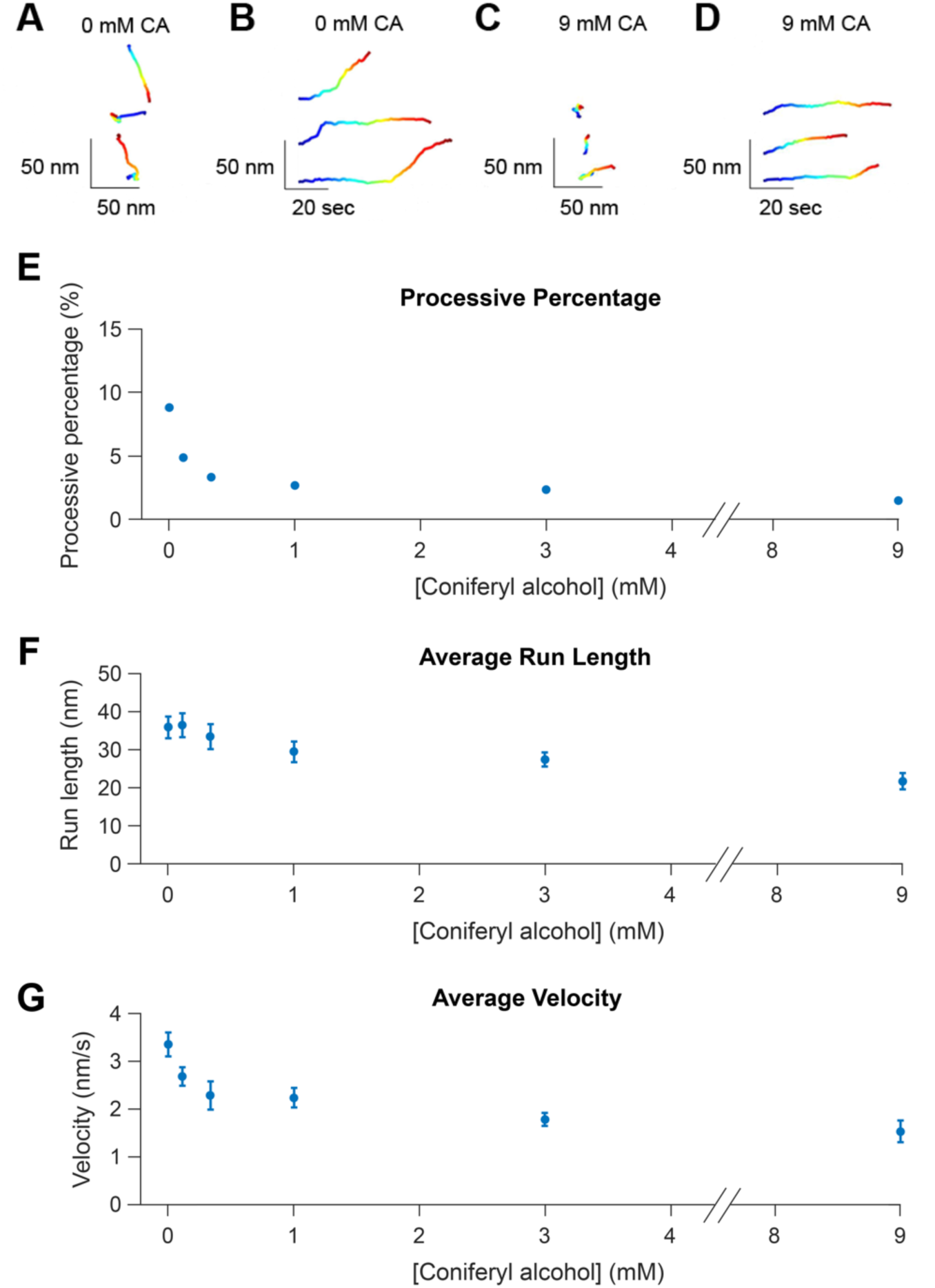
Cel7A molecules display diminishing processive behavior on cellulose with increasing lignification. Qdot-labeled Cel7A enzymes were imaged using total internal reflection microscopy at 1 frame/second and positions were fit using a Gaussian point-spread function. A: X-Y positions over time for three different Cel7A enzymes on lignin-free cellulose. B: Distance from origin versus time for the same three Cel7A enzymes on lignin-free cellulose shown in A. C: X-Y positions over time for three different Cel7A enzymes on lignocellulose prepared using 9 mM CA. D: Distance from origin versus time for the same three Cel7A enzymes on lignocellulose prepared using 9 mM CA shown in C. For all traces, blue represents the start of the binding event and red indicates the end of the binding event. E: Percentage of the total Cel7A enzymes imaged that displayed processive behavior. F: Average run length of processive segments; runs less than 10 nm long were excluded from analysis. G: Average velocity of processive segments; velocities less than 0.1 nm/s were excluded from analysis. Run lengths and velocities are presented as mean +/− SEM. The number of enzymes tracked for each sample were: 0 mM CA, N = 224; 0.11 mM CA, N = 172; 0.33 mM CA, N = 154; 1 mM CA, N = 93; 3 mM CA, N = 88; 9 mM CA, N = 56.

To characterize enzyme activity, we quantified the fraction of Cel7A molecules that moved processively, along with the velocities and run lengths of the processive moving enzymes. The percentage of processive molecules decreased with increasing amounts of lignin on the cellulose (Fig. 4E). In the cellulose-only samples, 8.8% of the molecules analyzed displayed processive behavior, consistent with previous work (33). With increasing levels of lignification, the processive fraction decreased sharply to a plateau at 1 mM CA, and at 9 mM CA the processive percentage was 1.5%, indicating an 83% reduction of the processive fraction. Run length also decreased from 36.0 ± 2.8 nm (mean ± SEM, N = 224 particles) in the cellulose-only sample to 21.7 ± 2.1 nm (mean ± SEM, N = 56 particles) on lignocellulose made with 9 mM CA, representing a 40% decline (Fig. 4F). The diminished run lengths of the processive enzymes suggest that at least part of the reduction in processive percentage can be explained by their run lengths falling below the 10 nm threshold we established for processive movement. Together, the shorter average run length and reduced fraction of processive enzymes are consistent with lignin acting as a barrier that impedes processive degradation of cellulose by Cel7A.

Finally, the Cel7A velocity, defined as the distance over the duration of processive segments, decreased by roughly half from 3.37 ± 0.25 nm/s (mean ± SEM, N = 224) in the control samples, consistent with previous work (33), to 1.54 ± 0.23 nm/s (mean ± SEM, N = 56) in the 9 mM CA samples (Fig. 4G). Similar to the processive percentage, the velocity fell steeply across the lower CA concentrations, where very little lignin was visible by electron microscopy (e.g., 0.11 mM CA in Fig. 2H). This slowing of Cel7A velocity indicates that, beyond acting as an impediment, lignin affects the ability of Cel7A to processively extract and hydrolyze the cellulose polymer.

### Lignin polymerized *in vitro* appears to form a thin layer on the cellulose surface

The decrease in the velocity and processive percentage at low CA concentrations where only sparse lignin labeling was observed by SEM raises the possibility that even in these samples there is a thin layer of lignin on the cellulose surface, invisible in SEM, that may slow the hydrolysis of cellulose by Cel7A. To investigate this possibility, we used fluorescence microscopy to test whether at low CA concentrations, lignin polymerizes on the cellulose surface in a form that is undetectable by IRM or SEM. To do this, we labeled lignin with Basic Fuchsin and measured the fluorescence intensity of the control and lightly lignified samples. Flow cells were formed with lignocellulose adhered to the surface of the glass coverslip, Basic Fuchsin was injected into the flow cell and incubated for 5 minutes, and excess dye was removed by extensively washing with sodium acetate buffer. Samples were then imaged using IRM to visualize the cellulose and by TIRFM to measure the fluorescence of Basic Fuchsin-stained lignin. 250-pixel by 250-pixel areas containing a minimal number of bright lignin aggregates were selected (yellow squares in Fig. 5D-F) and a histogram of lignin fluorescence intensities was generated from each sample. Similar labeling, illumination, and camera settings were used to enable quantitative comparison of lignin intensities across different samples.

**Figure 5:**
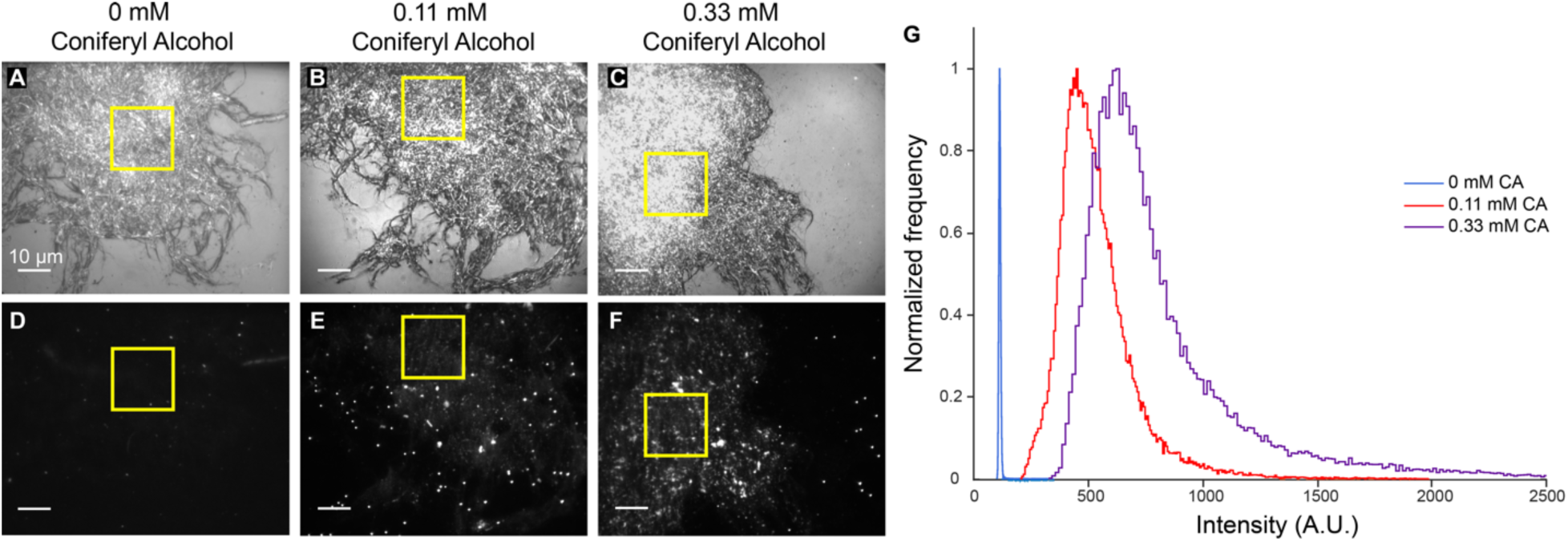
Lignin polymerized *in vitro* with low concentrations of coniferyl alcohol forms a thin layer of lignin that covers the cellulose surface. A-C: lignocellulose adhered to glass surface imaged using interference reflection microscopy; scale bar = 10 µm. D-F: total internal reflection fluorescence microscopy micrographs of lignocellulose stained with the lignin dye Basic Fuchsin; scale bar = 10 µm. G: Distribution of intensity values from Basic Fuchsin signal on lignocellulose surface of cellulose-only sample (blue), 0.11 mM CA lignocellulose sample (red), and 0.33 mM CA lignocellulose sample (purple). The frequency peaks for each sample were normalized to a value of 1 and the x-axis was truncated at 2500 A.U. Yellow squares indicate the 250-pixel by 250-pixel area of cellulose that intensity values were measured. Line scans that show fluorescence intensity across the samples are provided in Fig. S4.

In the cellulose-only sample, the Basic Fuchsin signal was negligible (Fig. 5D), with a very narrow distribution of intensities around a peak of 113 A.U. (Fig. 5G). This value is very close the background signal in cellulose-free regions of the coverslip (Fig. S4), indicating that Basic Fuchsin does not label cellulose. In the 0.11 mM and 0.33 mM CA samples, the lignin fluorescence increased across the cellulose surface as seen by eye in Figure 5E-F, and the pixel intensity distribution peaked at 448 A.U. and 621 A.U., respectively (Fig. 5G). The 0.33 mM CA sample contained lignin aggregates, visible by eye in the highlighted region, and these high intensities are apparent in the long tail of the 0.33 mM CA intensity distribution. Notably, there was virtually zero overlap in intensities between the cellulose-only control and the two lignified samples. This lack of overlap suggests that to the resolution of our fluorescence measurements, lignin is present everywhere on the cellulose surface, consistent with lignin forming a thin layer around the cellulose strands that is distinct from the small aggregates observed in the SEM and IRM images in Figure 2.2. This thin lignin layer provides a potential explanation for the slowing of Cel7A even at the lowest CA concentrations where deposition of obvious lignin aggregates is sparse.

## Discussion

In this work we investigated the potential mechanisms by which lignin inhibits degradation of lignocellulose by the cellulase Cel7A. Our data suggest that lignin does not act as a sink to irreversibly bind cellulases, but instead blocks Cel7A from binding to the cellulose surface, and acts as both an impediment to slow engaged enzymes and as a roadblock to block progress along a cellulose strand.

### *In vitro* polymerized lignin occludes the surface of cellulose

During the assembly of secondary plant cell walls, lignin is thought to polymerize onto cellulose-hemicellulose networks and form layers surrounding the microfibrils, with the degree of lignification depending on the type of biomass (41). While hemicellulose can be removed through various single-step pretreatments, G-lignin has proven to be difficult to fully remove from lignocellulosic biomass (42). High temperature alkaline pretreatment, one of the more effective approaches for lignin removal, alters the interaction between lignin and cellulose and causes lignin to form small aggregates (15, 27, 42–44). On the other hand, some acidic pretreatments cause the lignin to restructure around the cellulose into sheet-like structures (42).

Because we aimed to create lignocellulose samples that serve as a model for the lignocellulosic biomass used in biofuel production, we characterized our samples using SEM, IRM, and fluorescence microscopy. In the absence of lignin, the cellulose samples showed a meshwork of cellulose strands that intertwined with each other to create a complex web-like structure. This structure is roughly analogous to the bundled, multi-lamellate structure of cellulose in native plant cell walls (45), although cellulose in native walls is also typically interspersed in matrix polysaccharides such as hemicelluloses. Lignin polymerized in the absence of cellulose exhibited cauliflower-like assemblies under SEM, which appeared as high contrast ‘blobs’ in IRM. By SEM, lignin in the 0.11 mM and 0.33 mM CA samples appeared as small round aggregates in the cellulose meshwork. These structures may be analogous to the aggregates seen following high temperature alkaline pretreatment of native lignocellulose (15, 27, 42). Importantly, we observed Basic Fuchsin staining across the entire cellulose surface in these samples, suggesting that there is a thin coating of lignin across the surface that we do not detect by SEM or IRM. Evidence of lignin forming thin films in contact with cellulose has previously been reported (46, 47). By SEM, lignin in the 1 mM, 3 mM, and 9 mM CA samples appeared as larger and more structured assemblies on the cellulose surface and formed patches that wrapped around cellulose strands and occluded large regions of the cellulose surface (Fig. 2 and Fig. S1). These lignin structures may be analogous to the sheet-like structures seen following acid pretreatment of more highly lignified samples, such as softwood or wheat straw (27, 41).

### Cel7A does not accumulate on *in vitro* polymerized lignin

Published work has posited a ‘sink’ model in which lignin inhibits cellulase activity due to enzymes binding tightly and unproductively to lignin. One line of evidence comes from studies in which Cel7A is combined with cellulose or lignocellulose, the solution pelleted by centrifugation, and the amount of enzyme remaining in the supernatant compared. Diminished Cel7A in the supernatant for lignified cellulose samples was interpreted to mean that Cel7A binds directly to lignin (16, 22, 24, 25). A second line of evidence comes from experiments that measure the enzymatic activity of Cel7A by monitoring cellobiose production and find that activity is reduced in lignified cellulose compared to bare cellulose (13, 16, 22).

We found in fluorescence colocalization experiments (Fig. 3) that Cel7A does not accumulate on lignin and instead that regions of high lignin density correlate with less Cel7A binding in the presence of BSA. Without the addition of BSA in the samples, no correlation was measured between Cel7A binding and lignin deposition (Fig. S3). These data argue against the idea of lignin acting as a sink. Instead, they support a model of lignin blocking initial adsorption of Cel7A to cellulose by physically covering the cellulose surface. Further, they suggest that the decreased enzymatic activity of Cel7A due to lignin in bulk assays might result from lignin reducing the accessible surface area of cellulose, rather than from Cel7A irreversibly binding to lignin. Consistent with our results, cellulases show only modest binding to lignin in native cell walls in high-resolution microscopy experiments (27). One significant discrepancy between our experiments and previous work in bulk assays is that bulk assays use ∼10^3^ to 10^4^-fold higher enzyme concentrations than the nM concentrations used in our single-molecule studies. Thus, the negative Pearson’s correlation we see between cellulose and lignin binding indicates that Cel7A binds more tightly to cellulose than it does to lignin, but it does not rule out some nonspecific binding of Cel7A to lignin at the high enzyme concentrations used in bulk assays. Consistent with this, measured dissociation constants for cellulases binding to purified lignin are orders of magnitudes higher than the enzyme concentrations used in our assays (24, 25). Additionally, lignin in native lignocellulosic biomass is generally comprised of a mixture of monolignols (42), and future work is warranted to investigate whether cellulases bind more tightly to forms of lignin other than our pure synthetic G-lignin.

### *In vitro* polymerized lignin impedes Cel7A motion on cellulose

It is easy to envision that lignin aggregates and patches on the cellulose surface (Fig. 2) would disrupt interactions between Cel7A and cellulose, but Cel7A would be expected to interact normally in the microns-scale areas of cellulose that do not contain these large lignin assemblies. Consistent with this, the Cel7A run length, which is only tens of nm on lignin-free cellulose, was not affected by the lignin in the 0.11 mM and 0.33 mM CA samples. However, both the processive percentage and the velocity decreased steeply from the cellulose-only sample to the 0.33 mM CA sample (Fig. 4). Based on our fluorescence imaging of lignin by Basic Fuchsin, we interpret this inhibition to be due to a thin layer of lignin that coats the cellulose surface and is invisible in IRM and SEM. If the layer of lignin is thin enough to be penetrated or bypassed by the enzyme, the cellulase would still be able to hydrolyze the cellulose, but it might require more time to extract a cellulose chain from the fibril surface or its movement along the cellulose surface might be impeded, both of which would result in a lower velocity.

The decline in the processive percentage of Cel7A molecules with increasing lignin content might result from the lignin layer preventing Cel7A from accessing free ends of the cellulose or from extracting a cellulose chain from the crystal lattice. This finding that lignin decreased the percentage of processive Cel7A molecules provides another possible explanation for the reduced activity of Cel7A on lignified cellulose observed in bulk studies – because these static enzymes are likely not processively degrading cellulose, increasing the fraction of static enzymes is expected to decrease the overall catalytic turnover rate of the population of enzymes as measured in bulk. This diminished activity due to a reduced processive percentage would add to the expected reduced activity due to shorter run lengths and slower velocities.

### Conclusions

Based on the results of this study, we posit two molecular mechanisms to explain the inhibition of Cel7A activity by lignin (Fig. 6). We first propose that lignin on the cellulose surface impairs Cel7A activity by blocking Cel7A binding to cellulose. Second, aggregates and patches of lignin cannot be hydrolyzed by cellulases and act as roadblocks on the cellulase surface that impede Cel7A movement. At higher degrees of lignification, the lignin entraps cellulose strands and occludes the cellulose entirely. Based on the observation that no Cel7A accumulates on lignin under the conditions tested here, we conclude that lignin does not act as a sink, and instead acts to shield cellulose from Cel7A, such that Cel7A tends to bind to areas of cellulose that are not obstructed by lignin. Based on our observations at low degrees of lignification, we hypothesize that thin films of lignin can cover the cellulose surface and both diminish Cel7A velocity and reduce the percentage of enzymes that undergo processive movement on cellulose. In conclusion, lignin on the cellulose surface has a greater impact on Cel7A hydrolysis while lignin freely floating in solution would be expected to have a negligible effect on hydrolysis. Bearing in mind these inhibition mechanisms, we advocate that the primary focus for increasing cellulase activity on lignocellulosic biomass should be to disrupt the interaction between lignin and cellulose instead of removing lignin from biomass processing mixtures.

**Figure 6:**
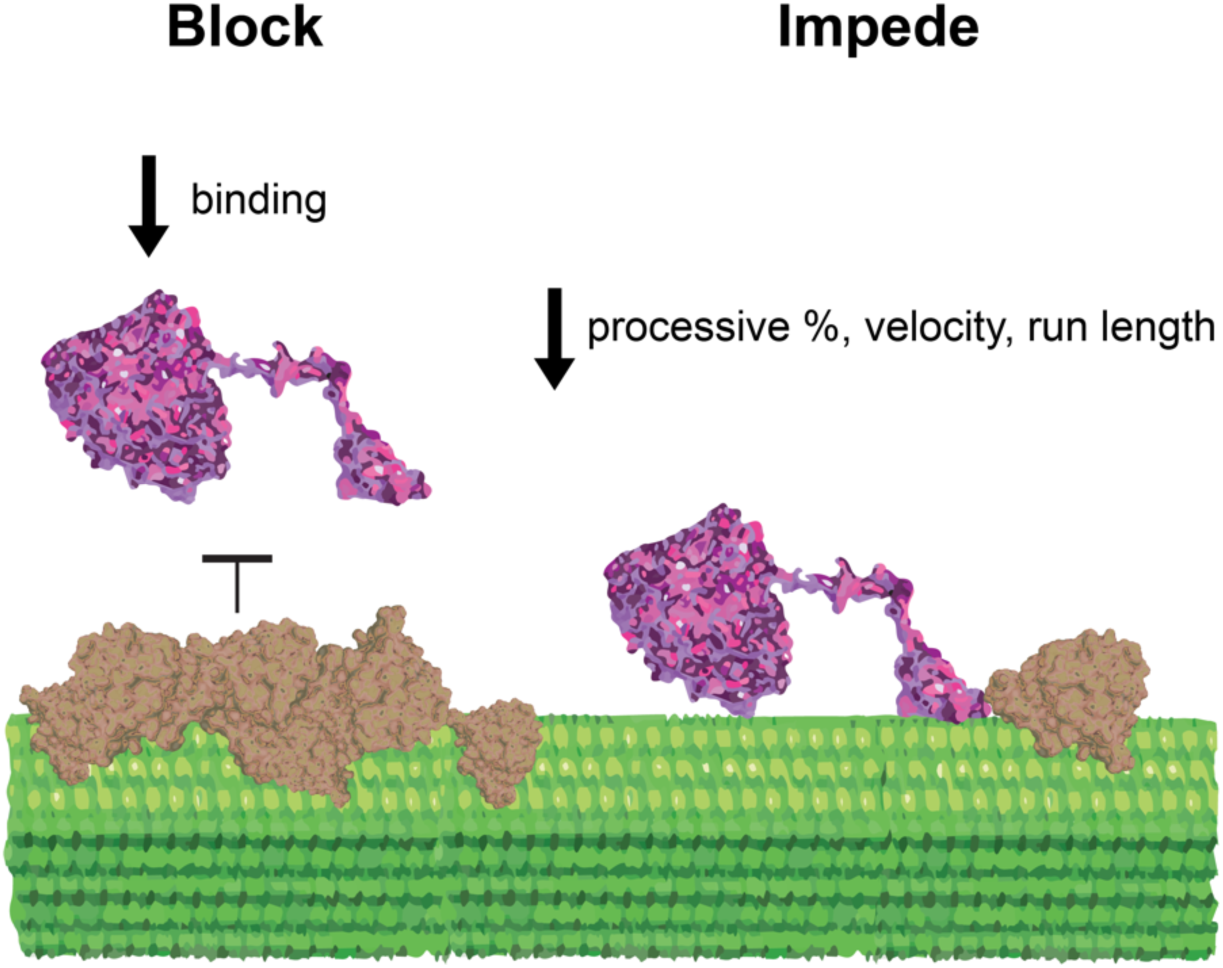
Models for Cel7A inhibition on cellulose caused by lignin as supported by single-molecule observations. Lignin covering the cellulose surface prevents Cel7A from binding to the cellulose substrate. Lignin on the cellulose surface can either partially or completely diminish the processive movement of Cel7A on cellulose. Lignin does not act as a sink due to the lack of Cel7A accumulation on lignin.

## Materials and Methods

### Cellulose and lignin preparation and characterization

Bacterial cellulose was produced by inoculating Schramm-Hestrin medium with *Gluconacetobacter hansenii* (strain ATCC 23769) (48), and growing the culture for 5 days at 30°C with no agitation. The resulting sheet of cellulose was washed five times with 100% ethanol. After filtration, 2% (w/v) NaOH was added to the cellulose and the solution was incubated for 30 minutes at 80°C. Next, the solution was centrifuged for 15 min at 2,300 rcf and the supernatant was decanted. The pellet was washed once with 0.5 M sodium acetate and twice with sterile ddH_2_O before being air dried for 2 days on aluminum foil. After drying, the cellulose was peeled off and stored at 4°C. Dried cellulose was re-suspended in 50 mL ddH_2_O and sonicated with a Sonic Dismembrator (Thermo Fisher, model 100) five times for 30 seconds each at a setting of 9, with 1 min breaks in between. Sonicated cellulose samples were combined and processed through a M-110EH microfluidizer at the Pennsylvania State University CSL Behring Fermentation Facility. The sample was first passed through a 200 µm filter five times at 5,000 psi and then passed through a 75 µm filter for 45 minutes at 7,000 psi. The cellulose content was determined by phenol sulfuric acid using a glucose standard (49).

Lignocellulose samples were created by combining cellulose with coniferyl alcohol (CA), hydrogen peroxide, and horseradish peroxidase (HRP) to polymerize G-lignin on the cellulose surface (34). Two different methods were used to produce the lignin, the first used increasing (HRP) catalyst concentrations proportionally with the hydrogen peroxide and CA concentrations, and the second maintained a constant HRP concentration while the hydrogen peroxide and CA concentrations were increased. Based on the SEM and single-molecule results, both methods of lignin polymerization showed similar effects, so the data were combined into a single data set. Lignin was polymerized by adding 0.11 mM – 9 mM CA and H_2_O_2_ to 0.5 mL of 4.25 mM acetobacter cellulose along with either 0.01 mg/mL – 0.9 mg/mL HRP for 1 hour at 37°C or 0.01 mg/mL HRP for 20 hours at 25°C. The lignocellulose samples were then centrifuged at 9,000 rcf for 5 minutes and gently resuspended in ddH_2_O so the final concentration of cellulose was 4.5 mM. The removed supernatant was analyzed by measuring the light absorbance at 260 nm to confirm that the oxidation reaction of CA went to completion.

### Cel7A preparation and characterization

*T. reesei* Cellobiohydrolase I (Sigma-Aldrich, catalog number: E6412), hereafter referred to as Cel7A, was buffer exchanged into 50 mM sodium acetate using a PD-10 column (General Electric). Peak fractions, as determined by absorbance at 280 nm, were pooled and glycerol was added for a final concentration of 10% (v/v). The final protein concentration (6.02 μM) was determined by light absorbance, using an extinction coefficient of 74,906 M^−1^cm^−1^. Cel7A was biotinylated using EZ-Link NHS-LC-LC-Biotin (Thermo Scientific, catalog number: 21343), which labels the primary amines of exposed lysine residues. Cel7A was buffer exchanged into 50 mM NaBO_3_ (pH 8.5), combined with biotin-NHS dissolved in dried Dimethylformamide (DMF) at a biotin:enzyme ratio of 10:1, and incubated for 3 hours in the dark at 21°C. To remove the free biotin, the enzymes were buffer exchanged back into 50 mM sodium acetate using a PD-10 desalting column. The enzyme concentration was calculated using absorbance measurements at 280 nm and the biotin concentration was determined using the Pierce Fluorescence Biotin Quantitation Kit (Thermo Scientific, catalog number: 46610). The biotin:Cel7A ratio was determined to be 0.60. Biotinylated enzymes were flash frozen using liquid nitrogen and stored at −80°C. After thawing for experiments, enzymes were never refrozen.

### Single-molecule TIRFM imaging and analysis

To prepare flow cells, a ∼10 µL volume of 4.5 mM lignified acetobacter cellulose was pipetted onto the surface of a glass slide, as previously described (33). Two strips of double-sided tape were positioned on either side of the cellulose solution and a plasma cleaned glass cover slip was placed on top of the tape to create a flow cell (∼30 µL volume). The slide was inverted and placed into an oven at 65°C for 30 minutes to allow the cellulose solution to dry, leaving the cellulose fibers stuck to the surface of the cover slip. TetraSpeck beads (Thermo Scientific; catalog number: T7280) used as fiduciary markers were then injected into the flow cell and incubated for 5 minutes to allow them to non-specifically bind to the glass surface. This was followed by three washes of 1 mg/mL bovine serum albumin (BSA) with three minutes incubation each, to prevent nonspecific binding of cellulase enzymes to the glass surface.

Qdot-labeled Cel7A was prepared by mixing 3 nM biotinylated Cel7A with 2 nM Qdot 655 (Thermo Scientific; catalog number: Q10123MP) in 50 mM sodium acetate, pH 5.0, with 5 mM dithiothreitol to prevent photobleaching. Following a 15-minute incubation, the solution was injected into the flow cell. Decreasing the enzyme:particle ratio below this led to many fewer landing events, consistent with Qdots binding single enzymes. Single-molecule imaging was accomplished using total internal reflection fluorescence microscopy (TIRFM) with an excitation laser of 488 nm at 30 mW power to illuminate both the TetraSpeck beads on the surface and the Qdots attached to the enzymes (32). Cellulose was visualized by interference reflectance microscopy (IRM) with a white light LED. Images were taken at 1 frame/s and videos consisted of 1,000 frames. The imaging area for each frame was 79.2 μm x 79.2 μm with a pixel size of 73 nm per pixel. A quadrant photodiode (QPD) sensor connected to the microscope stage prevented drift in the z-direction to keep the images in constant focus. All videos were captured at 21°C.

ImageJ was used to combine two 500-frame videos captured consecutively of the same region of interest to create the final 1,000 frame videos. Videos were analyzed using FIESTA software, which fits two-dimensional Gaussians to the point spread functions of the TetraSpeck beads and the Qdot-labeled cellulases to create single-molecule trajectories (40). The resulting traces were imported into scripts written in MATLAB for further analysis of individual tracks, as described in previous works (32, 33). The positional changes of TetraSpeck beads were subtracted from all tracks to correct for stage drift in the X-Y direction. Particles with total binding durations of less than 10 seconds were not included in the analysis because it was often difficult to differentiate processive segments from spatial variances observed in static segments. Few molecules had binding durations greater than 510 seconds, but those that did were excluded from the analysis, as they were potentially due to irreversible binding by denatured enzymes, and thus were considered outliers.

### Scanning electron microscopy imaging and analysis

Lignocellulose samples were washed six times for 5 minutes each using a Millipore 0.2 µm membrane filter starting at 50% ethanol and increasing the ethanol concentration to 60%, 70%, 85%, 95%, and 100% ethanol. A Leica EM CPD300 was used for critical point drying of the lignocellulose on the membrane filter. The dried samples were then mounted onto an aluminum stub with carbon tape and stored in a desiccator at room temperature until the day of experimentation. The samples were sputter coated with ∼10 nm of gold-palladium and visualized using a Zeiss SIGMA VP-FESEM at the Pennsylvania State University Huck Institutes of Life Sciences. Images were captured at 90,000x magnification with a EHT of 3 kV using type II secondary electrons.

## Supplemental Figures

**Figure S1 (Related to Figure 2):**
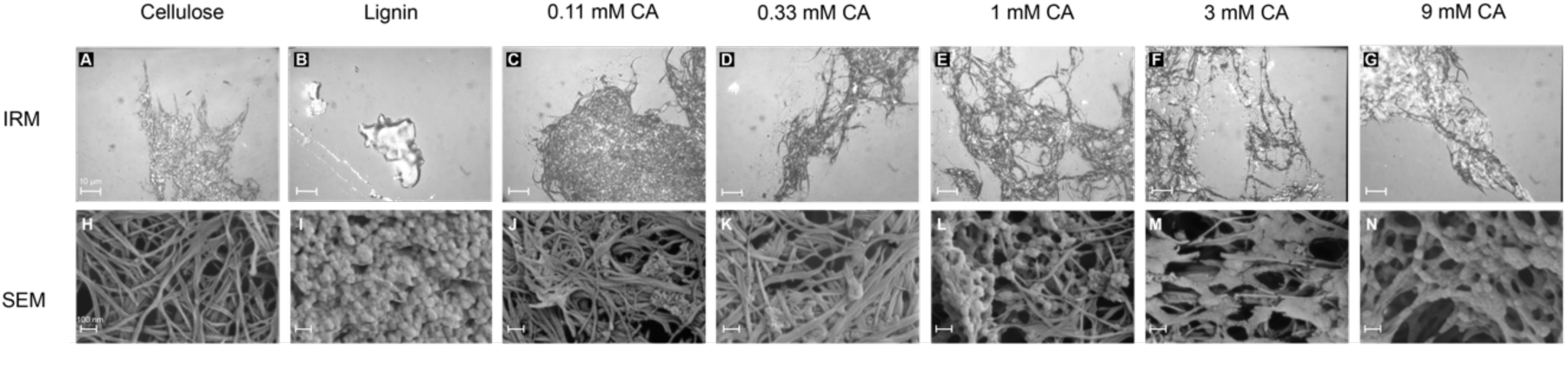
All CA concentrations used in data set for lignin polymerized onto acetobacter cellulose *in vitro*. First column: cellulose only; second column: lignin only. Third through fifth columns: cellulose with lignin polymerized from different concentrations of CA. Top row are interference reflection micrographs, scale bar = 10 µm. Bottom row are scanning electron micrographs, scale bar = 100 nm.

**Figure S2 (Related to Figure 2):**
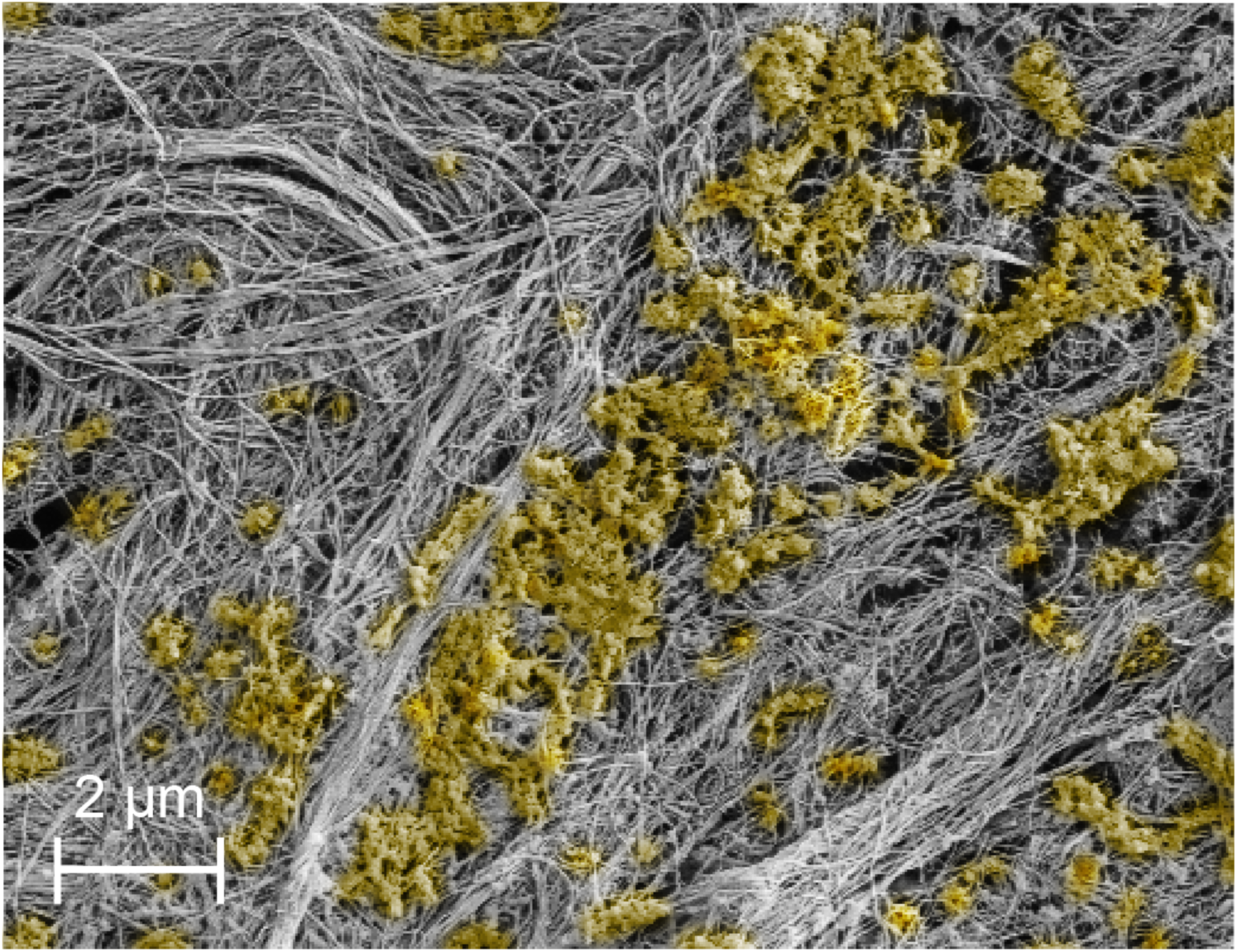
Lignin generated *in vitro* deposits heterogeneously onto acetobacter cellulose. Scanning electron micrograph of 3 mM CA sample, with lignin false-colored yellow. Areas in the middle of the image display highly lignified regions of cellulose with sheets of lignin (shown in yellow) covering significant areas of the cellulose surface In contrast, areas in the upper left and bottom right of the image contain less lignin on the cellulose surface, with some regions appearing to have nearly no visible lignin and appearing similar to the cellulose-only samples.

**Figure S3 (Related to Figure 3):**
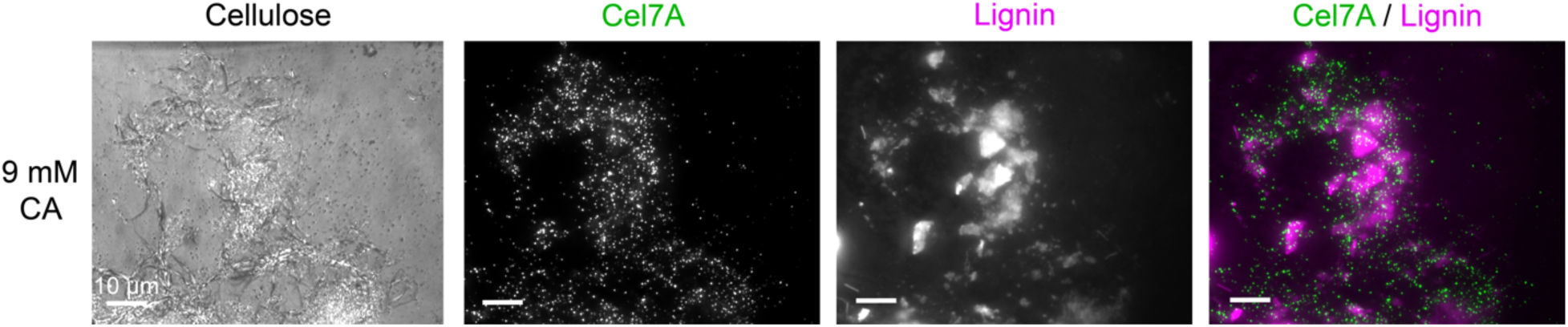
BSA reduces the binding of Cel7A to lignin. 9 mM CA lignocellulose was adsorbed to the slide without the addition of BSA to the flow cell to determine if BSA affects Cel7A binding to lignin. Binding locations of Qdot-labeled Cel7A were recorded for 50 seconds before being washed out and Basic Fuchsin was added to determine locations of lignin deposition. A Pearson’s correlation coefficient of 0.037 was calculated, as described in Methods. This value indicates no correlation between Cel7A binding and lignin, which differs from previous results with BSA added to the flow cell as shown in Figure 3. This shows the BSA washes may affect the Cel7A binding to lignin, but the lignin still does not appear to act as a sink where a larger positive correlation would be expected.

**Figure S4 (Related to Figure 5):**
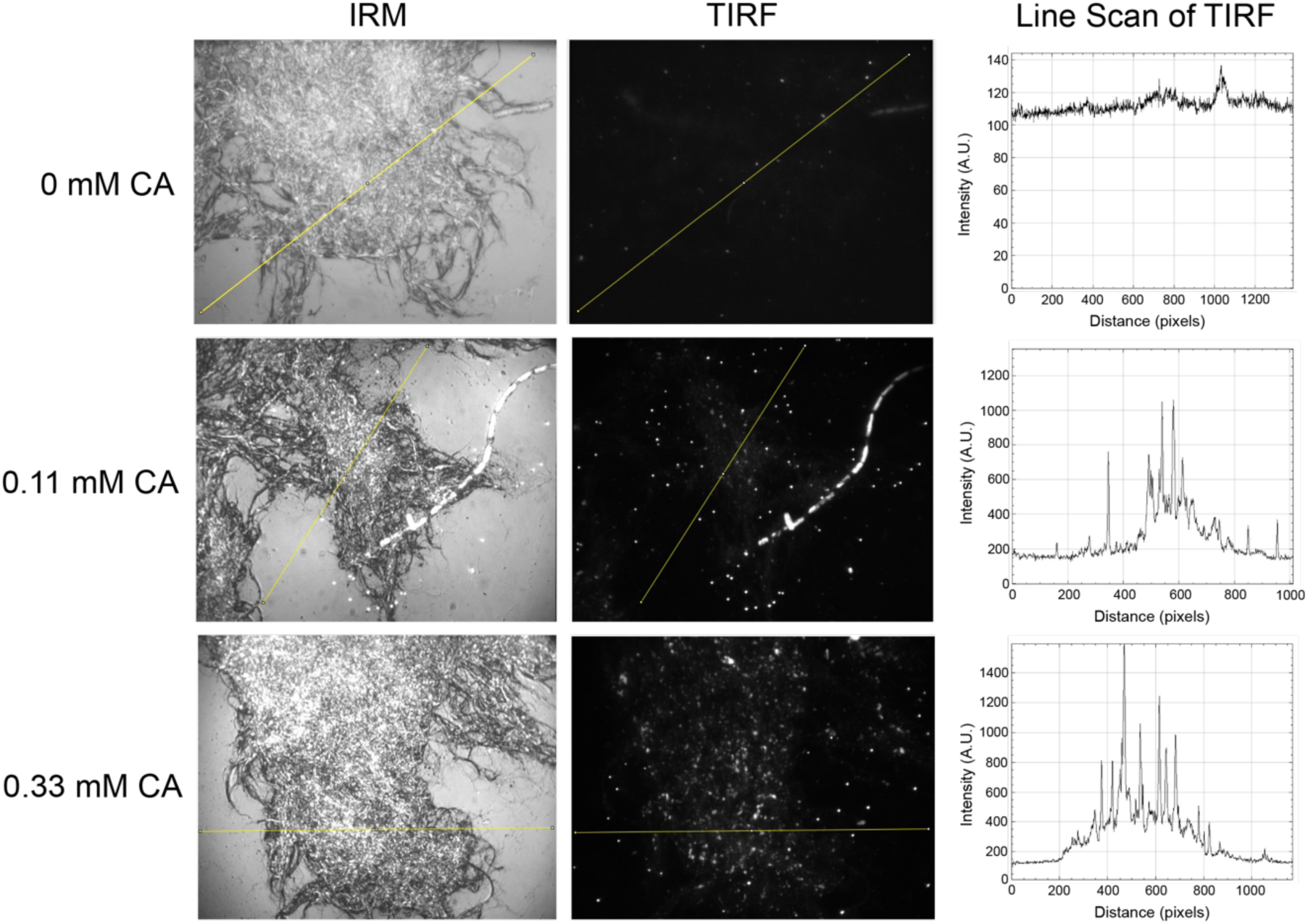
Line scans across the lignocellulose surfaces show an increase in Basic Fuchsin fluorescent signal in the TIRF channels on the lignified cellulose samples compared to cellulose-only samples. Yellow lines in the IRM and TIRF images show the location of the line scan for each sample. The fluorescence intensity across the cellulose surface for the cellulose-only sample is similar to the intensity on the glass surface, indicating no lignin is present in the sample. The 0.11 mM and 0.33 mM CA samples display an increase in fluorescence intensity across the lignocellulose surface compared to the glass surface, signifying the presence of a thin film of lignin on the cellulose surface. The line scans for the lignified samples also show periodic spikes, corresponding to lignin aggregates that can be seen by eye in the fluorescence image.

